# Prediction of brain age in individuals with and at risk for alcohol use disorder using brain morphological features

**DOI:** 10.1101/2024.03.01.582844

**Authors:** Chella Kamarajan, Babak A. Ardekani, Ashwini K. Pandey, Jacquelyn L. Meyers, David B. Chorlian, Sivan Kinreich, Gayathri Pandey, Christian Richard, Stacey Saenz de Viteri, Weipeng Kuang, Bernice Porjesz

## Abstract

Brain age measures predicted from structural and functional brain features are increasingly being used to understand brain integrity, disorders, and health. While there is a vast literature showing aberrations in both structural and functional brain measures in individuals with and at risk for alcohol use disorder (AUD), few studies have investigated brain age in these groups. The current study examines brain age measures predicted using brain morphological features, such as cortical thickness and brain volume, in individuals with a lifetime diagnosis of AUD as well as in those at higher risk to develop AUD from families with multiple members affected with AUD (i.e., higher family history density (FHD) scores). The AUD dataset included a group of 30 adult males (mean age = 41.25 years) with a lifetime diagnosis of AUD and currently abstinent and a group of 30 male controls (mean age = 27.24 years) without any history of AUD. A second dataset of young adults who were categorized based on their FHD scores comprised a group of 40 individuals (20 males) with high FHD of AUD (mean age = 25.33 years) and a group of 31 individuals (18 males) with low FHD (mean age = 25.47 years). Brain age was predicted using 187 brain morphological features of cortical thickness and brain volume in an XGBoost regression model; a bias-correction procedure was applied to the predicted brain age. Results showed that both AUD and high FHD individuals showed an increase of 1.70 and 0.09 years (1.08 months), respectively, in their brain age relative to their chronological age, suggesting accelerated brain aging in AUD and risk for AUD. Increased brain age was associated with poor performance on neurocognitive tests of executive functioning in both AUD and high FHD individuals, indicating that brain age can also serve as a proxy for cognitive functioning and brain health. These findings on brain aging in these groups may have important implications for the prevention and treatment of AUD and ensuing cognitive decline.

## 1. Introduction

Prediction of brain age relative to one’s chronological age can serve as a useful biomarker of brain health [1,2]. Brain age estimation using neural measures has been successfully used as a measure of neurobiological aging and cognitive decline/lag [3–5] in clinical populations [1,2,6,7]. Brain age indices computed from structural and functional brain measures, especially using brain morphological features, are increasingly being used to understand brain integrity, neurodevelopment, and neuropsychiatric disorders [2,8–10]. There is substantial evidence that substance use disorders can lead to accelerated biological aging [11], including brain aging [12,13] and cognitive impairment [14–16].

Alcohol misuse is a leading cause for premature death and disability [17,18]. While there is a vast literature showing aberrations in both structural and functional brain measures in individuals with and at risk for alcohol use disorder (AUD) [19,20], few studies have investigated brain age in these groups using neurophysiological features. There is strong neuroimaging evidence that chronic AUD can lead to accelerated aging of brain morphology and hypothesized contribution to age-related dementia [21]. It is also well-established that children from families densely affected with AUD families who are at higher risk to develop AUD, manifest neurocognitive deficits and atypical structural and functional brain features [22–24]. Therefore, it is important to measure and understand changes in brain age in individuals with AUD as well as high-risk individuals from high-dense AUD families.

Neuroimaging-based methods are predominantly used to estimate brain age and *brain age gap (BAG),* a measure to quantify the deviation of an individual from age-dependent distributions that are meant to model normal aging [2,25]. Brain age measures are usually calculated by first modeling the association between selected brain-derived measurements and chronological age in a healthy (or control) sample to serve as a baseline, and then the trained model will be applied to a sample of interest (usually the clinical group) to predict brain age at the individual level [2]. Recently, there have been accumulating efforts to develop and test methods to measure brain age using machine learning models [2,8,9,26–29]. More recently, several studies have used XGBoost [30], a popular, efficient ensemble based algorithm [8,9], has been frequently used in recent studies to predict brain age [8,9,13], as it has been shown to have superior predictive power and model performance compared to other machine learning methods [29]. It is important to note that predicted brain age often involves bias in the estimation as the computed score is overestimated in younger individuals and underestimated in older individuals in the initial prediction before any correction [31–34], and therefore, bias-correction procedures are in place [35]. For the current study, we will use the correction procedure adopted in a recent work by de Lange et al. [8], who have demonstrated brain age prediction predicted age using neuroanatomical features in an XGBoost model.

The current study has estimated brain age in two datasets related to alcohol use phenotypes: (i) a group of adults with a lifetime diagnosis of AUD along with a comparison group of healthy controls, and (ii) a group of young adults at higher risk to develop AUD from families with multiple members affected with AUD (i.e., higher family history density (FHD) of AUD scores) and a low-risk comparison group with relatively low FHD scores. We have used XGBoost regression model to predict brain age in both datasets. We have also implemented age-bias correction in order to optimize brain age measurement. As most of the recent studies have used MRI-based neuroanatomical features to predict brain age [1,5–8,27,36–38], we have specifically used cortical thickness and volumetric measures following a recent study by Rutherford et al. [27]. In order to understand the association between brain age and cognitive performance, we have included neuropsychological measures of executive functions and memory performance in the study. We expected accelerated brain age in the AUD group as well as in the high-risk group with high FHD scores compared to those with low FHD scores. The present study can enhance our understanding of brain aging in individuals with AUD as well as in high-risk individuals with high scores of family history density of AUD.

## 2. Materials and Methods

### 2.1. Participants

The sample consisted of two different datasets: (i) The AUD dataset comprised a total of 60 male participants (age range = 19.75–51.08 years), in which 30 individuals had a lifetime diagnosis of DSM-IV alcohol dependence (AUD group; M_age_=41.25; SD_age_=7.20; age range = 25.58–51.08 years) and 30 individuals did not have any AUD diagnosis (CTL group; M_age_=27.24; SD_age_=4.78; age range = 19.75–38.08 years) [see **Table 1**]. The AUD individuals were recruited from alcoholism treatment centers in and around NYC after they had been detoxified and did not have any withdrawal or other major AUD symptoms during the time of scanning. CTL individuals were recruited through advertisements and screened to exclude any personal and/or family history of major medical, psychiatric, or substance-related disorders. A detailed description of this dataset is provided in our previous publications [39,40]. (ii) The FHD dataset had a total of 71 participants (38 males and 33 females), of whom 40 individuals (20 males) were grouped as *HiFHD* based on their high scores (above the median value of 5) on the measure of family history density (FHD) [41], while the remaining 31 individuals (18 males) were categorized as *LoFHD* as they had low FHD scores (below the median value of 5) [see **Table 2**]. FHD scores, which ranged from 0 to 1, were computed as the proportion of non-descendant First and Second-degree relatives with DSM-5 diagnosis of Alcohol Use Disorder; a detailed description of this measure is available in our previous publication by Pandey et al. [41].

**Table 1:** Comparison of clinical features between the AUD and control participants.

| Variables | AUD |  |  | Control |  |  | t | p |
| --- | --- | --- | --- | --- | --- | --- | --- | --- |
|  | N<sup>a</sup> | Mean | SD | Na | Mean | SD |  |  |
| Age (in years) | 30 | 41.25 | 7.20 | 30 | 27.24 | 4.78 | 8.78 | <0.0001 |
| Alcohol: Age of onset (regular use)<sup>b</sup> | 30 | 15.77 | 2.58 | 12 | 20.5 | 3.8 | -4.67 | <0.0001 |
| Alcohol: Quantity/day (heavy use period)<sup>c</sup> | 30 | 11 | 7.66 | 12 | 3 | 1.6 | 5.58 | <0.0001 |
| Alcohol: Frequency/month (heavy use period)<sup>c</sup> | 30 | 20 | 9.01 | 12 | 4 | 4.09 | 7.97 | <0.0001 |
| Alcohol: Quantity/day (last 6 months)<sup>c</sup> | 30 | 3 | 6.61 | 18 | 3 | 1.98 | 0.045 | NS |
| Alcohol: Frequency/month (last 6 months)<sup>c</sup> | 30 | 4 | 8.02 | 18 | 3 | 3.62 | 0.6 | NS |
| Length of Abstinence (in days)<sup>d</sup> | 30 | 672.93 | 844.94 | 18 | 57 | 149.76 | 3.89 | <0.0005 |
| Tobacco: Quantity/day (last 6 months)<sup>c</sup> | 20 | 10 | 5.8 | 6 | 2 | 1.63 | 5.19 | <0.0001 |
| Tobacco: Frequency/month (last 6 months)<sup>c</sup> | 20 | 28 | 4.83 | 6 | 14 | 13.82 | 2.47 | NS |
| Marijuana: Frequency (last 6 months)<sup>c</sup> | 10 | 99 | 91.38 | 4 | 19 | 27.61 | 2.58 | <0.03 |

<sup>a</sup> Note that all participants were assessed but the data of participants who did not report on specific items were omitted from the analysis here.
<sup>b</sup> Drink one day/per month for 6 months or more.
<sup>c</sup> Quantity and Frequency mean values were rounded to the nearest whole numbers as per the reviewers' suggestion.
<sup>d</sup> Range (5–4228 days).

**Table 2:** Comparison of substance use features between the HiFHD and LoFHD groups.

| Substance | Category | HiFHD |  |  | LoFHD |  |  | t-value | p-value |
| --- | --- | --- | --- | --- | --- | --- | --- | --- | --- |
|  |  | N | Mean | SD | N | Mean | SD |  |  |
| Age | Demography | 40 | 25.33 | 3.74 | 31 | 25.47 | 3.40 | 0.1435 | 0.4430 |
| Alcohol | Dependence Dx | 40 | 0.03 | 0.16 | 31 | 0.10 | 0.30 | 1.2980 | 0.1990 |
|  | Dependence Sx | 40 | 0.78 | 1.37 | 31 | 0.94 | 1.12 | 0.5290 | 0.5980 |
|  | Abuse Dx | 40 | 0.13 | 0.34 | 31 | 0.06 | 0.25 | -0.8400 | 0.4040 |
|  | Abuse Sx | 40 | 0.30 | 0.76 | 31 | 0.16 | 0.45 | -0.9000 | 0.3710 |
| Tobacco | Dependence Dx | 40 | 0.15 | 0.36 | 31 | 0.10 | 0.30 | -0.6610 | 0.5110 |
|  | Dependence Sx | 40 | 0.85 | 1.64 | 31 | 0.97 | 1.60 | 0.3030 | 0.7630 |
| Marijuana | Dependence Dx | 40 | 0.53 | 1.45 | 31 | 0.16 | 0.37 | -1.3600 | 0.1780 |
|  | Dependence Sx | 40 | 1.45 | 2.01 | 31 | 1.19 | 1.70 | -0.5690 | 0.5710 |
|  | Abuse Dx | 40 | 0.38 | 1.44 | 31 | 0.06 | 0.25 | -1.1810 | 0.2420 |
|  | Abuse Sx | 40 | 0.48 | 0.82 | 31 | 0.26 | 0.63 | -1.2230 | 0.2250 |
| Cocaine | Dependence Dx | 40 | 0.05 | 0.22 | 31 | 0.00 | 0.00 | -1.2590 | 0.2120 |
|  | Dependence Sx | 40 | 0.28 | 1.26 | 31 | 0.03 | 0.18 | -1.0620 | 0.2920 |
|  | Abuse Dx | 40 | 0.00 | 0.00 | 31 | 0.00 | 0.00 | NA | NA |
|  | Abuse Sx | 40 | 0.13 | 0.56 | 31 | 0.00 | 0.00 | -1.2340 | 0.2220 |
| Opioid | Dependence Dx | 40 | 0.03 | 0.16 | 31 | 0.00 | 0.00 | -0.8790 | 0.3830 |
|  | Dependence Sx | 40 | 0.13 | 0.79 | 31 | 0.06 | 0.36 | -0.3950 | 0.6940 |
|  | Abuse Dx | 40 | 0.00 | 0.00 | 31 | 0.00 | 0.00 | NA | NA |
|  | Abuse Sx | 40 | 0.03 | 0.16 | 31 | 0.00 | 0.00 | -0.8790 | 0.3830 |
| Stimulant | Dependence Dx | 40 | 0.00 | 0.00 | 31 | 0.00 | 0.00 | NA | NA |
|  | Dependence Sx | 40 | 0.00 | 0.00 | 31 | 0.00 | 0.00 | NA | NA |
|  | Abuse Dx | 40 | 0.00 | 0.00 | 31 | 0.00 | 0.00 | NA | NA |
|  | Abuse Sx | 40 | 0.00 | 0.00 | 31 | 0.00 | 0.00 | NA | NA |
| Sedative | Dependence Dx | 40 | 0.00 | 0.00 | 31 | 0.03 | 0.18 | 1.1380 | 0.2590 |
|  | Dependence Sx | 40 | 0.05 | 0.32 | 31 | 0.19 | 1.08 | 0.8010 | 0.4260 |
|  | Abuse Dx | 40 | 0.00 | 0.00 | 31 | 0.00 | 0.00 | NA | NA |
|  | Abuse Sx | 40 | 0.00 | 0.00 | 31 | 0.10 | 0.54 | 1.1380 | 0.2590 |

Dx – Diagnosis, Sx – Symptom Counts, NA – Not Applicable due to zero values

A modified version of the semi-structured assessment for the genetics of alcoholism (SSAGA) [42], a polydiagnostic clinical interview was administered to assess alcohol/substance use and related disorders. Participants were instructed to abstain from alcohol and other substances for at least 5 days prior to the scanning. Standard MRI safety protocols and exclusion criteria (implants, tattoos, cosmetics, claustrophobia, etc.) were followed to ensure subjects’ safety and data quality. Individuals with hearing/visual impairment, a history of head injury, or moderate and severe cognitive deficits (<21) on mini-mental state examination (MMSE) [43] were also excluded from the study. All participants provided informed consent and the Institutional Review Board approved the respective study protocols.

### 2.2. MRI Data Acquisition and Image Processing

The MRI data were collected using three different systems: (i) AUD dataset (N=60) was collected with Siemens 3T Tim *Trio*; (ii) first batch of 59 individuals from the FHD dataset were scanned with Siemens 3T Magnetom *Skyra*; and (iii) second batch of 12 participants from the FHD dataset were scanned with Siemens 3T Magnetom *Prisma*. A high-resolution three-dimensional T1-weighted magnetization-prepared rapid gradient-echo (MPRAGE) image was collected with the following parameters for each scanner: (i) *Trio*: TR=2500 ms, TE=3.5 ms, TI=1200 ms, flip angle=8°, matrix size=256×256×192, and voxel size=1×1×1 mm^3^; (ii) *Skyra*: TR = 2100 ms, TE = 4.78 ms, TI = 900 ms, flip angle = 8°, matrix size = 256×256×176, and voxel size = 1×1.2×1.2 mm^3^; and (iii) *Prisma*: TR=2300 ms, TE=2.98 ms, TI=900 ms, flip angle=8°, matrix size=240×256×160, and voxel size=1×1×1 mm^3^. Standard procedures and protocols of MRI scanning were followed.

A total of 187 brain morphological features [see **Table A1** in the *Appendix section*] were extracted for the prediction of brain age, based on the previous study by Rutherford et al. [27]. We used the *sMRIPprep* (version 0.12.2) workflow [44], which is part of a Python-based image analysis platform *Nipype* (version 1.8.6) [45], for preprocessing anatomical data from the T1- weighted (T1w) images. Complete details of the software, processing, processing pipeline and documentation are available at https://www.nipreps.org/smriprep/. First, the T1-weighted (T1w) image was corrected for intensity non-uniformity (INU) with *N4BiasFieldCorrection* [46], distributed with *ANTs* (version 2.3.3) [47], and used as T1w-reference throughout the workflow. The T1w-reference was then skull-stripped with a Nipype implementation of the *antsBrainExtraction.sh* workflow (from ANTs), using *OASIS30ANTs* as target template. Brain tissue segmentation of cerebrospinal fluid (CSF), white-matter (WM) and gray-matter (GM) was performed on the brain-extracted T1w using *fast* from FSL (version 6.0.5.1:57b01774) [48].

Brain surfaces were reconstructed using *recon-all* from FreeSurfer (version 7.3.2) [49], and the brain mask estimated previously was refined with a custom variation of the method to reconcile ANTs-derived and FreeSurfer-derived segmentations of the cortical gray-matter of *Mindboggle* [50]. Cortical thickness values of the Destrieux parcellation [51] containing 74 regions and a mean value for each hemisphere, along with volumetric measure for 37 brain structures, were extracted from the Freesurfer output directories. All 187 neuroanatomical features (150 on cortical thickness and 37 on brain volumes) were used in the brain age prediction model.

### 2.3. Brain Age Prediction

We implemented an *XGBoost* [30] (stands for *eXtreme Gradient Boosting*) as several recent studies have used this algorithm to predict brain age using MRI [52–57] as well as EEG measures [29]. The details of the algorithm and computational steps are available in the work by de Lange et al. [8], wherein the algorithms from the native XGBoost [30] and Scikit-learn [58] have been implemented for computing brain age. As mentioned earlier, 187 anatomical features (i.e., 150 features of bilateral cortical thickness and 37 features of brain volumes) [see **Table A1** in *Appendix section*] served as the predictor variables against chronological age as the response or outcome variable in the XGBoos regression model. The prediction models were run separately for the AUD and FHD datasets. We trained the model using the reference groups (i.e., CTL group for the AUD dataset and the LoFHD group for the FHD dataset), while tested the prediction model on the case groups (AUD group for the AUD dataset and the HiFHD group for the FHD dataset), as it is usually done in the brain age studies [2]. Predicted brain age was subjected to an age-bias correction procedure [8,35]. While training the model, the values of the predictor variables (i.e., the neuroanatomical features) were scaled using the *robust scaler* [59] from the *scikit-learn* library [58], which removes the median and scales the data according to the quantile range [8]. Hyper parameter tuning was implemented in order to obtain the best model parameters to use in the training phase based on the previous work [8]. We tuned the model parameters using inner 3-fold and outer 5-fold cross-validation procedure, following the work by de Lange et al. [8]. The *SearchTerm* space parameters in the model were: *max_depth* = range(1, 11, 2), *n_estimators* = range(50, 400, 50), *learning_rate* = [0.001, 0.01, 0.1, 0.2]. According to this procedure, a correction is applied to the predictions by first fitting *Y = α × Ω + β*, where Y is the modelled predicted age as a function of chronological age (Ω), and α and β represent the slope and intercept [8]. The derived values of α and β are then used to correct predicted age with *Corrected Predicted Age = Predicted Age + [Ω − (α × Ω + β)].* The brain age gap, a measure of lag or difference between the chronological age and the predicted brain age [8,60], was also computed. The model performance was measured using the correlation coefficient (r), r-square (R^2^), Mean Absolute Error (MAE), and Root Mean Squared Error (RMSE). In this study, the correlation coefficient (r) between the chronological age and corrected predicted age served as the primary model performance. Higher r-value (regardless of the direction) indicated better performance. While R^2^, which is not the square of r in this case, is a measure of predictive power of the regression in terms of how much variation is explained by the regression, and therefore higher R^2^ represents better performance. The MAE represents the average of the absolute value of each residual, and lower MAE will imply better performance.

On the other hand, the RMSE is computed as the square root of the average of squared errors, where the errors are squared before they are averaged. Similar to MAE, higher RMSE signifies lower performance. The MAE and the RMSE together are useful to estimate the variation in the errors in a set of predictions. The RMSE is larger or equal to the MAE and difference between the two measures reflects the magnitude of the variance in the individual errors in the sample.

### 2.4. Neuropsychological Assessment

Participants in both datasets were administered two computerized tests from the Colorado assessment tests for cognitive and neuropsychological assessment [61], namely, the Tower of London Test (TOL) [62], and the visual span test (VST) [63]. Details are available in our previous publication [64].

The TOL assesses the planning and problem-solving ability of the executive functions. In this test, participants were asked to solve a set of puzzles with graded difficulty levels by arranging a specific number of colored beads one at a time from a starting position to the desired goal position in as few moves as possible. The test consisted of 3 puzzle types, with 3, 4, and 5 colored beads placed on the same number of pegs, with 7 problems/trials per type and a total of 21 trials. The following performance measures from the sum total of all puzzle types were used in the analysis: (i) actual moves made (MovMade), (ii) excess moves, which is the additional moves beyond the minimum moves required to solve the puzzle (“ExcMovMade”); (iii) percentage of excess moves (PctAbvOpt), (iv) average pickup time, which is the initial thinking/planning time spent until picking up the first bead to solve the puzzle (“AvgPicTime”); (v) average total time, which is the total thinking/planning time to solve the problem in each puzzle type (“AvgTotTime”); (vi) total trial time, which is the total performance/execution time spent on all trials within each puzzle type (“TotTrlTime”); and (vii) average trial time, which is the mean performance/execution time across trials per puzzle type (“AvgTrlTime”).

The VST measured visuospatial memory span from the forward condition and working memory from the backward condition. In this test, a set of randomly arranged squares, ranging from 2 to 8 squares per trial, flashed in a predetermined sequence depending on the span level being assessed. Each span level was administered twice, with a total of 14 trials in each condition. During the forward condition, subjects were required to repeat the sequence in the same order by clicking on the squares using a computer mouse. In the backward condition, subjects were required to repeat the sequence in the reverse order (starting from the last square). The following performance measures were collected during forward and backward conditions: (i) total number of correctly performed trials (“TotCor_Fw” and “TotCor_Bw”); (ii) maximum span or sequence-length achieved (“Span_Fw” and “Span_Bw”); (iii) total average time, which is the sum of mean time-taken across all trials performed (“TotAvgTime_Fw” and “TotAvgTime_Bw”); and (iv) total correct average time, which is the sum of mean time-taken across all trials correctly performed (“TotCorAvgTime_Fw” and “TotCorAvgTime_Bw”).

### 2.5. Statistical Analysis

Statistical analyses were done using the SPSS software package (Version 28.0.1.1). We compared the demographic and clinical characteristics across the groups of AUD and FHD datasets respectively using independent samples t-tests. Pearson correlations were used to find association among all age variables (i.e., chronological age, uncorrected brain age, corrected brain age, uncorrected brain age gap and corrected brain age gap) as well as between specific age measures and neuropsychological and clinical measures. Distributions of some of the brain age measures were plotted for visualization.

## 3. Results

### 3.1. Comparison of Clinical Variables Across the Groups

As shown in Table 1, abstinent AUD participants were significantly younger than control participants. AUD participants started drinking earlier than controls. During their past heavy use period, AUD participants drank more in terms of quantity and frequency than control participants who endorsed drinking. However, the quantity and frequency of drinking during the past 6 months were not significantly different between the AUD and control participants. At the time of assessments, the AUD individuals were maintaining abstinence from drinking longer than the control participants who endorsed drinking. Tobacco use was significantly higher among the AUD participants during the past 6 months than the control participants who endorsed tobacco use, although the frequency of use was not statistically significant. Marijuana use during the past months was also higher among AUD participants compared to the controls.

### 3.2. Brain Age Prediction in the AUD Dataset

In the XGBoost regression model with AUD dataset, the best model parameters were the following: n_estimators=200, max_depth=9, and learning_rate=0.01. The prediction performance during the training phase in terms of the absolute mean (SD) values for R^2^, MAE, and RMSE are: 0.096 (0.096), 3.814 (1.828), and 4.443 (1.672), respectively. **Fig. 1** shows the distribution of the individual values of Corrected Brain Age against Chronological Age (left panels) and Corrected Brain Age Gap (right panels) for the Control group (top panels) and the AUD group (bottom panels). It can be observed that a majority of the AUD individuals (bottom left panel) show increased brain age (shown over the referenced diagonal line).

**Fig. 1.**
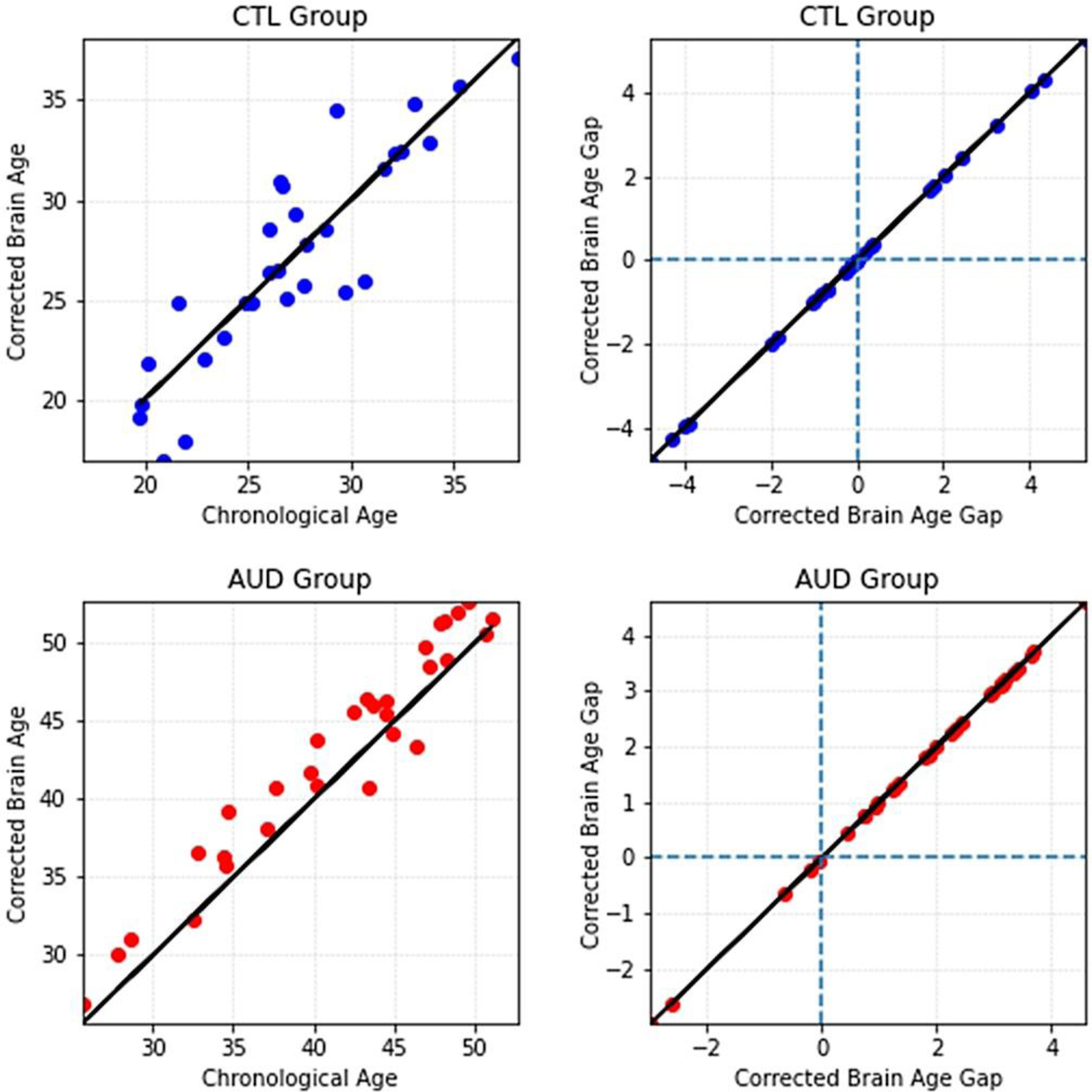
Distribution of the individual values of Corrected Brain Age against Chronological Age (left panels) and Corrected Brain Age Gap (right panels) for the Control group (top panels) and the AUD group (bottom panels). The individual values (marked in dots) above the diagonal line in the left panels represent increased brain age while the values below the diagonal line represent decreased brain age.

Bar Graphs in **Fig. 2** show the mean values for all age measures in the AUD dataset for the age measures (Chronological Age, Uncorrected Brain Age, and Corrected Brain Age) [top panel] as well as for the brain age gap measures (Uncorrected Brain Age Gap and Corrected Brain Age Gap) [bottom panel] for the AUD dataset. It is shown that the Uncorrected Brain Age (29.01 years) in the AUD group was possibly underpredicted by the model (before age-bias correction) against the Chronological Age (41.25 years) during the testing phase. On the other hand, the Corrected Brain Age (42.95 years) was 1.70 years higher than the Chronological Age (41.25 years) as illustrated in the bar graphs. Control group, which was the training dataset), did not show visible changes in the brain ages. The same trend is seen in terms of the Brain Age Gap (i.e., the difference between chronological age and the brain age) measures shown in the bottom panel in **Fig. 2**.

**Fig. 2.**
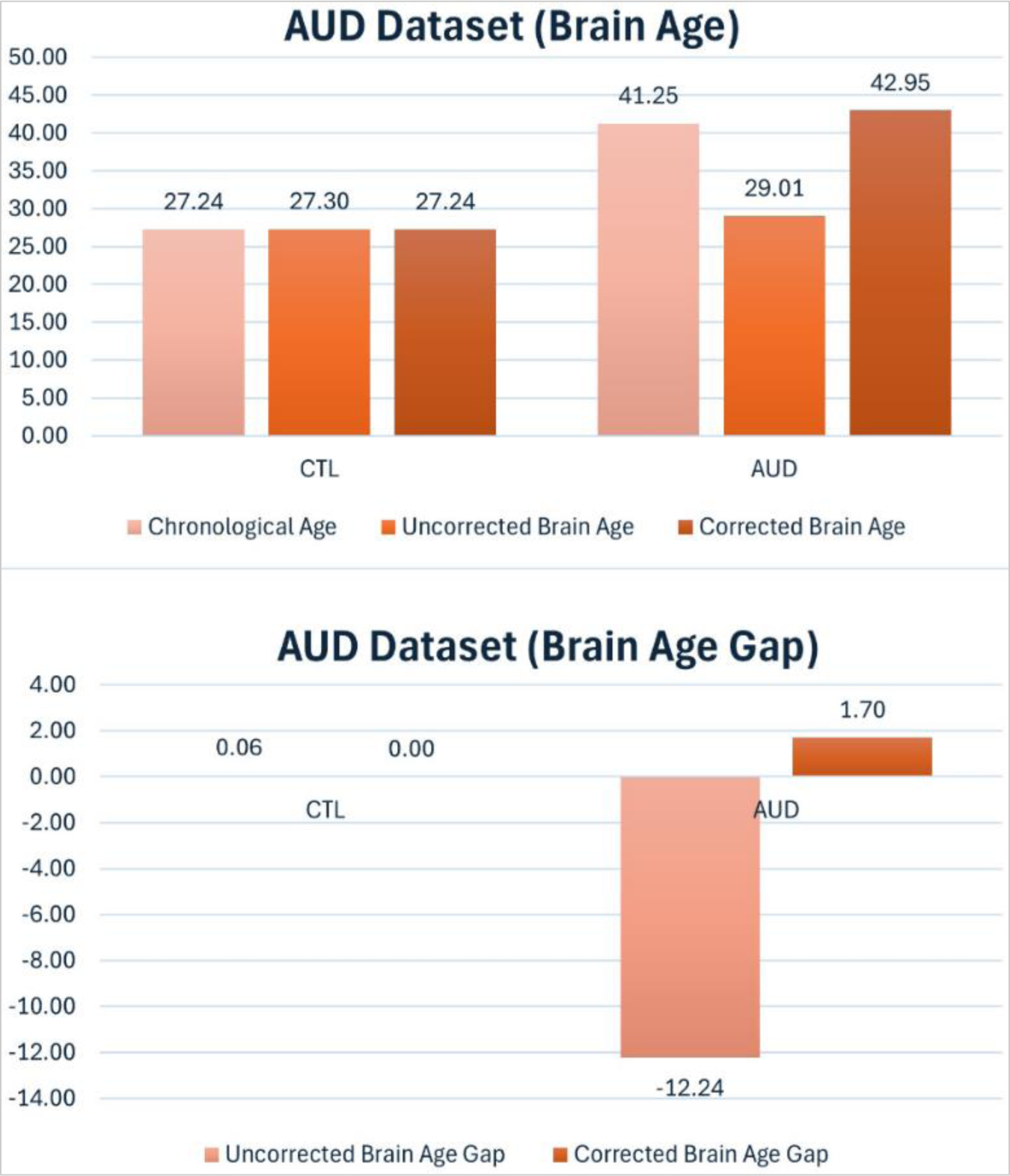
Bar Graphs showing the mean values for Chronological Age, Uncorrected Brain Age, and Corrected Brain Age (top panel) as well as for Uncorrected Brain Age Gap and Corrected Brain Age Gap (bottom panel) for the AUD dataset.

### 3.3. Brain Age Prediction in the FHD Dataset

In the XGBoost regression model with FHD dataset, the best model parameters were the following: n_estimators=300, max_depth=9, and learning_rate=0.001. The prediction performance in terms of the absolute mean (SD) values for R^2^, MAE, and RMSE are: 0.704 (0.985), 2.822 (0.697), and 3.431 (0.959), respectively. **Fig. 3** shows the distribution of the individual values of Corrected Brain Age against Chronological Age (left panels) and Corrected Brain Age Gap (right panels) for the Lo-FHD group (top panels) and the Hi-FHD group (bottom panels). It can be observed that a considerable portion of the HiFHD individuals (bottom left panel), especially after the age 22, show increased brain age relative to their respective chronological age.

**Fig. 3.**
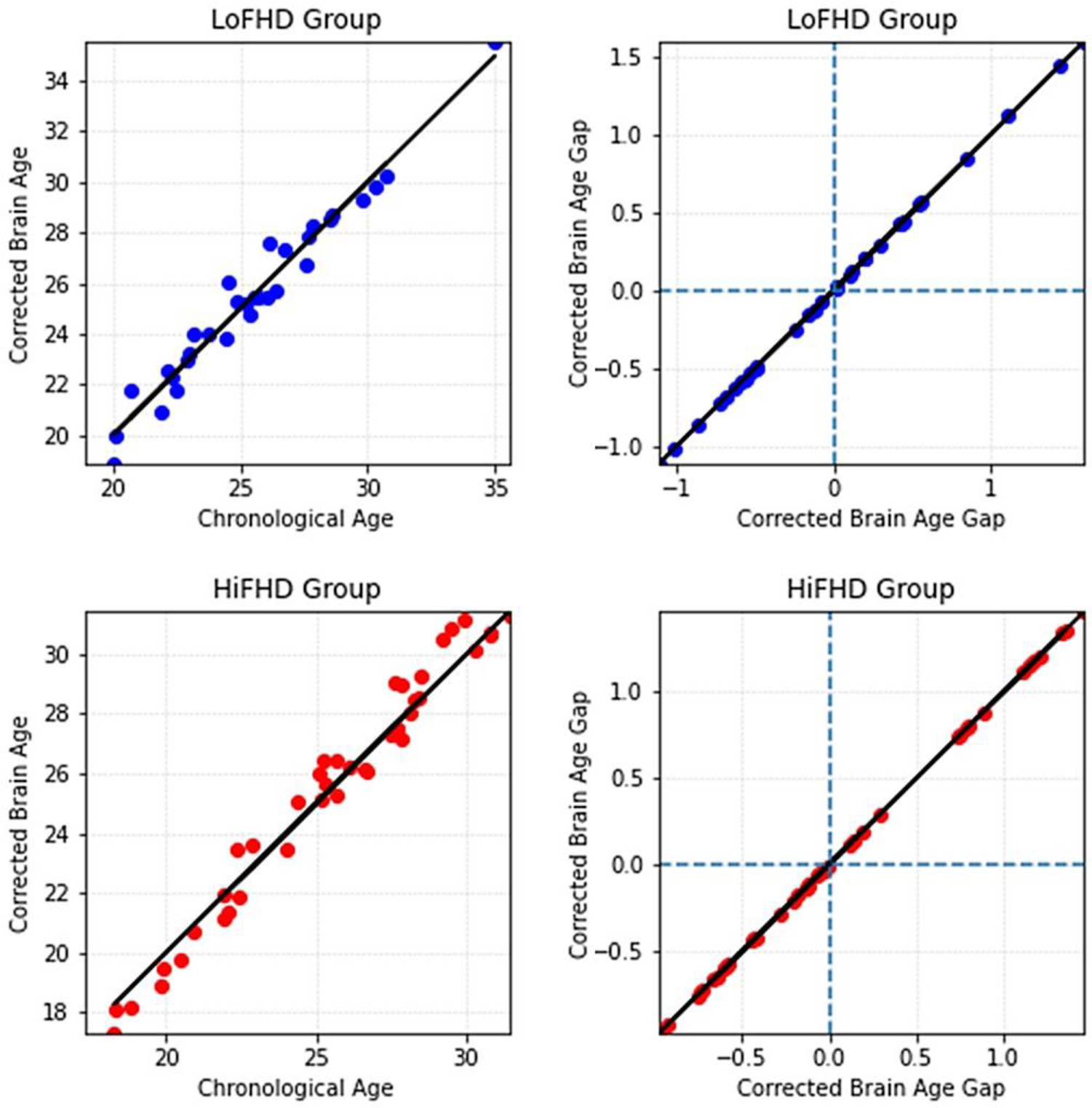
Distribution of the individual values of Corrected Brain Age against Chronological Age (left panels) and Corrected Brain Age Gap (right panels) for the LoFHD group (top panels) and the HiFHD group (bottom panels). The individual values (marked in dots) above the diagonal line in the left panels represent increased brain age while the values below the diagonal line represent decreased brain age.

Bar Graphs in **Fig. 4** show the mean values for all age measures in the AUD dataset for the age measures (Chronological Age, Uncorrected Brain Age, and Corrected Brain Age) [top panel] as well as for the brain age gap measures (Uncorrected Brain Age Gap and Corrected Brain Age Gap) [bottom panel] for the AUD dataset. These graphs show that the Uncorrected Brain Age in the HiFHD group was the highest followed by the Corrected Brain Age (during the testing phase). The Corrected Brain Age (25.42 years), which is the measure of our interest, was 0.09 years (1.08 months) higher than the Chronological Age (25.33 years). Interestingly, the brain age starts to steadily increase from age 22 until early 30s in this group, and the mean difference between Chronological Age and Corrected Brain Age in this subgroup (age ≥22; N=30) rises to 0.2334 years (2.80 months) [see **Fig. 3**, bottom left panel]. On the other hand, the LoFHD group did not show any visible changes in the Corrected Brain Age (during the training phase), as expected.

**Fig. 4.**
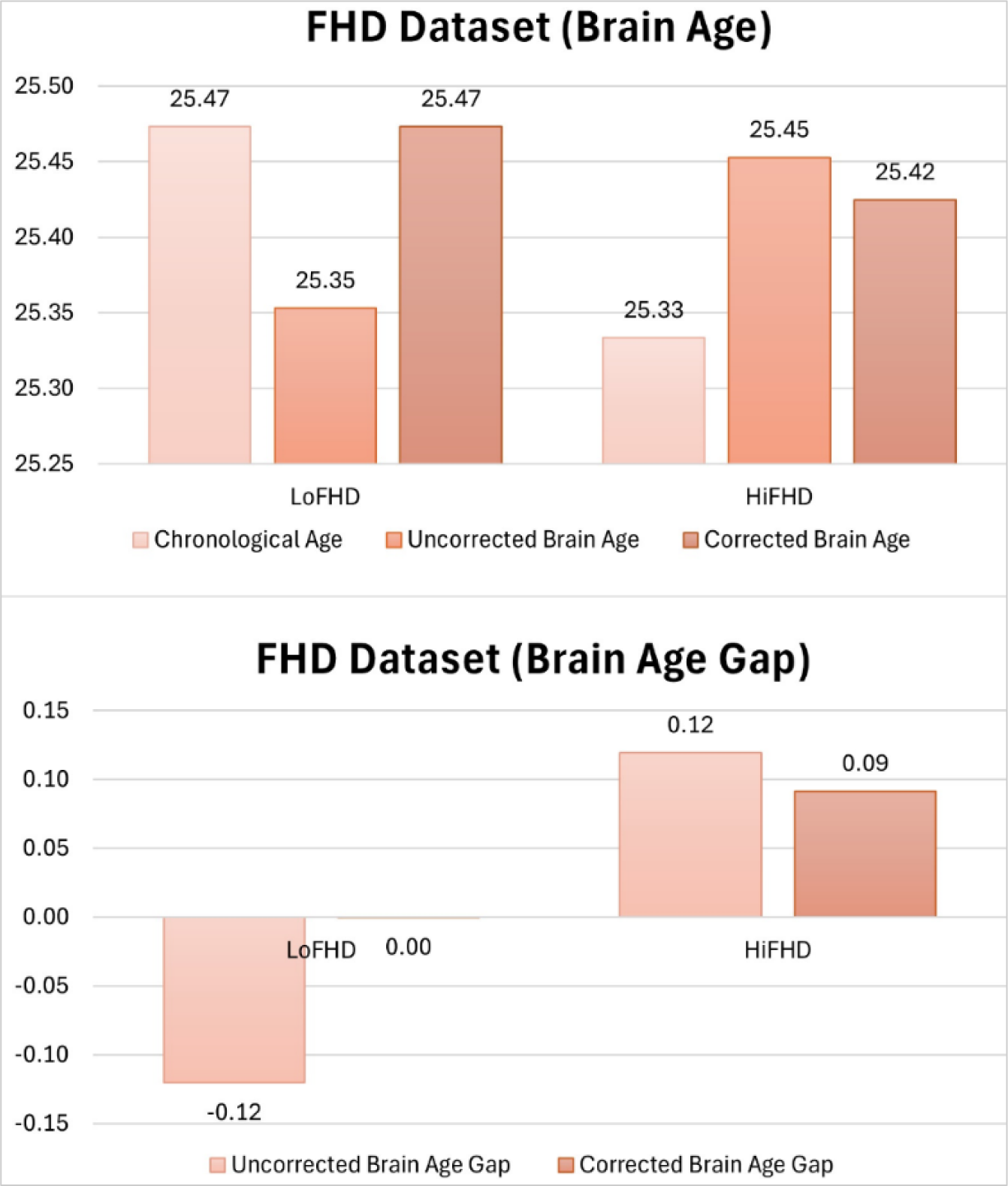
Bar Graphs showing the mean values for Chronological Age, Corrected Brain Age, and Uncorrected Brain Age (top panel) as well as for Brain Age Gap (bottom panel) for the FHD dataset.

### 3.4. Correlations of Age Measures with Themselves and Other Variables

#### 3.4.1. Correlations across the Age Measures

Pearson correlations among the brain age measures in the AUD dataset are shown in **Table 3**. In the Control group, significant correlations were observed between (1) Chronological Age and Uncorrected Brain Age, (2) Chronological Age and Uncorrected Brain Age Gap, (3) Uncorrected Brain Age and Corrected Brain Age, (4) Uncorrected Brain Age and Uncorrected Brain Age Gap, (5) Uncorrected Brain Age and Corrected Brain Age Gap, (6) Corrected Brain Age and Uncorrected Brain Age Gap, (7) Corrected Brain Age and Corrected Brain Age Gap, and (8) Uncorrected Brain Age Gap and Corrected Brain Age Gap. On the other hand, the AUD group showed significant correlations between (1) Chronological Age and Corrected Brain Age, (2) Chronological Age and Uncorrected Brain Age Gap, (3) Uncorrected Brain Age and Uncorrected Brain Age Gap, (4) Uncorrected Brain Age and Corrected Brain Age Gap, (5) Corrected Brain Age and Uncorrected Brain Age Gap, and (6) Uncorrected Brain Age Gap and Corrected Brain Age Gap.

**Table 3:** Pearson correlations among the brain age measures in the AUD dataset. The correlation values in the lower triangle represent the control group and those in the upper triangle represent the AUD group.

|  |  | AUD |  |  |  |  |
| --- | --- | --- | --- | --- | --- | --- |
|  |  | Chronological Age | Uncorrected Brain Age | Corrected Brain Age | Uncorrected Brain Age Gap | Corrected Brain Age Gap |
| CTL | Chronological Age | - | -0.1457 | 0.9691*** | -0.9731*** | -0.1502 |
|  | Uncorrected Brain Age | 0.0022 | - | 0.1031 | 0.3696* | 1.0000*** |
|  | Corrected Brain Age | 0.8903** | 0.4574* | - | -0.8861*** | 0.0985 |
|  | Uncorrected Brain Age Gap | -0.8900** | 0.4539* | -0.5847** | - | 0.3739* |
|  | Corrected Brain Age Gap | 0 | 1.0000** | 0.4555* | 0.4559* | - |

\*\*\*p<0.001; \*\*p<0.01; \*p<0.05

Pearson correlations among the brain age measures in the FHD dataset are shown in **Table 4**. The Low-FHD (LoFHD) group showed significant correlations between (1) Chronological Age and Uncorrected Brain Age, (2) Chronological Age and Uncorrected Brain Age Gap, (3) Uncorrected Brain Age and Uncorrected Brain Age Gap, (4) Uncorrected Brain Age and Corrected Brain Age Gap, and (5) Corrected Brain Age and Uncorrected Brain Age

**Table 4:** Pearson correlations among the brain age measures in the FHD dataset. The correlation values in the lower triangle represent the Low-FHD (LoFHD) group and those in the upper triangle represent the High-FHD (HiFHD) group.

|  |  | Hi-FHD |  |  |  |  |
| --- | --- | --- | --- | --- | --- | --- |
|  |  | Chronological Age | Uncorrected Brain Age | Corrected Brain Age | Uncorrected Brain Age Gap | Corrected Brain Age Gap |
| **Lo-FHD** | Chronological Age | - | 0.1431 | 0.9875*** | -0.9846*** | 0.4387** |
|  | Uncorrected Brain Age | -0.2847 | - | 0.2975 | 0.0323 | 0.9521*** |
|  | Corrected Brain Age | 0.9805*** | -0.0907 | - | -0.9446** | 0.5750*** |
|  | Uncorrected Brain Age Gap | -0.9826*** | 0.4580** | -0.9268*** | - | -0.2747 |
|  | Corrected Brain Age Gap | 0 | 0.9586*** | 0.1966 | 0.1859 | - |

\*\*\*p<0.001; \*\*p<0.01; \*p<0.05

Gap. On the other hand, significant correlations in the High-FHD (HiFHD) group were observed between (1) Chronological Age and Corrected Brain Age, (2) Chronological Age and Uncorrected Brain Age Gap, (3) Chronological Age and Corrected Brain Age Gap, (4)

Uncorrected Brain Age and Corrected Brain Age Gap, (5) Corrected Brain Age and Uncorrected Brain Age Gap, and (6) Corrected Brain Age and Corrected Brain Age Gap.

#### 3.4.2. Correlations of Age Measures with Neuropsychological Performance

Correlations of specific age measures with neuropsychological scores from the Tower of London (TOL) Test and the Visual Span Test (VST) in the AUD dataset are shown in **Table 6**. It

**Table 6:** Correlations of specific age measures with neuropsychological scores from the Tower of London (TOL) Test and the Visual Span Test (VST) in the AUD dataset.

| Neuropsychological Variables<sup>ψ</sup> | AUD |  | CTL |  |
| --- | --- | --- | --- | --- |
|  | Chronological Age | Corrected Brain Age | Chronological Age | Corrected Brain Age |
| TOL_MovMade | 0.4909* | 0.4924* | 0.0236 | 0.0535 |
| TOL_ExcMovMade | 0.4909* | 0.4924* | 0.0236 | 0.0535 |
| TOL_PctAbvOpt | 0.4909* | 0.4924* | 0.0235 | 0.0535 |
| TOL_AvgPicTime | 0.0761 | 0.0334 | 0.1849 | 0.2519 |
| TOL_AvgTotTime | 0.1555 | 0.0901 | 0.1759 | 0.2296 |
| TOL_TotTrlTime | 0.3801 | 0.3399 | 0.1843 | 0.2374 |
| TOL_AvgTrlTime | 0.3801 | 0.3399 | 0.1843 | 0.2374 |
| VST_TotCor_Fw | -0.4275* | -0.3956 | -0.0183 | -0.0254 |
| VST_Span_Fw | -0.4147* | -0.3906 | -0.0339 | -0.0265 |
| VST_TotAvgTime_Fw | 0.0037 | -0.0366 | 0.1010 | 0.1133 |
| VST_TotCorAvgTime_Fw | -0.0101 | -0.0514 | 0.1400 | 0.1322 |
| VST_TotCor_Bw | -0.2895 | -0.2549 | -0.1274 | -0.0815 |
| VST_Span_Bw | -0.2859 | -0.2628 | -0.0717 | -0.0245 |
| VST_TotAvgTime_Bw | -0.0793 | -0.1233 | -0.0228 | -0.0206 |
| VST_TotCorAvgTime_Bw | -0.0548 | -0.1297 | -0.0993 | -0.0226 |

\*p<0.01
<sup>ψ</sup>Refer Section 2.4. for the details of the neuropsychological variables listed in this table.

is observed that three of the TOL neuropsychological variables related to number of moves made (i.e., total moves, excess moves, and percentage above optimal moves) showed significant positive correlations with both chronological age and corrected brain age but only in the AUD group, suggesting poorer performance in executive functioning (planning) with increasing chronological and brain ages. Further, two of the VST neuropsychological variables related to forward trials (i.e., total correct and maximum span) significant negative correlations with chronological age in the AUD group, suggesting poorer memory performance with advancing age. There were no significant correlations observed in the CTL group participants.

In the FHD dataset (**Table 7**), two of the TOL neuropsychological variables on performance time (i.e., average pickup time, and average total performance time) showed significant positive correlations with both chronological age and corrected brain age but only in the High FHD group, suggesting poor neuropsychological performance with increasing age. None of the VST variables were significant. There were no significant correlations observed in the CTL group participants.

**Table 7:**
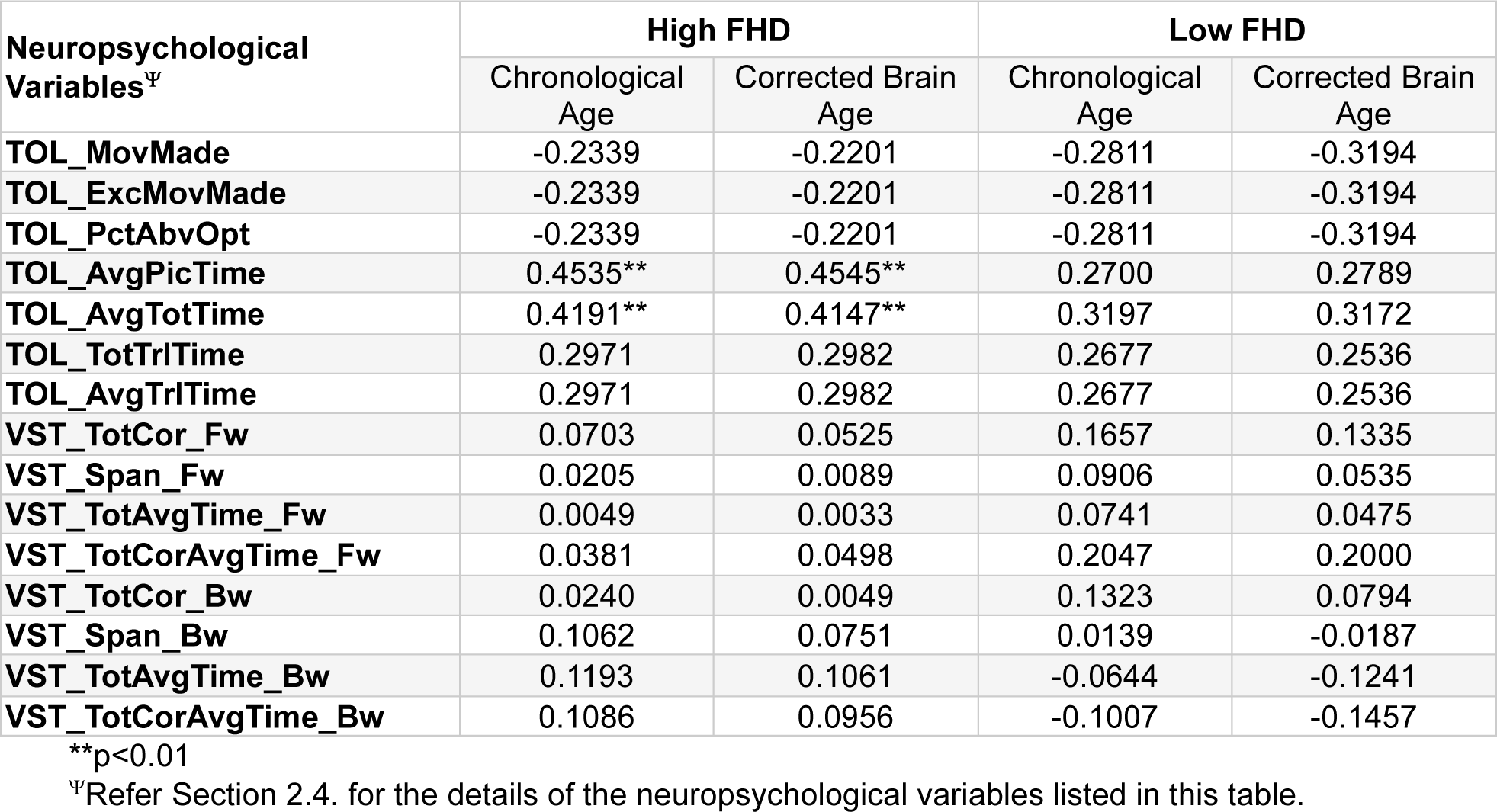
Correlations of specific age measures with neuropsychological scores from the Tower of London (TOL) Test and the Visual Span Test (VST) in the FHD dataset.

## 4. Discussion

The objective of the present study was to predict brain age using neuroanatomical features such as cortical thickness and brain volume in individuals with a lifetime diagnosis of AUD as well as in those with increased family history density (FHD) in their first- and degree relatives.

Results showed that both AUD and high-risk individuals showed an increase of 1.70 and 0.09 years (1.08 months), respectively, in their brain age relative to their chronological age, suggesting accelerated brain aging in these individuals. Increased brain age was found to be associated with poor neuropsychological performance. The significance and implications of the major findings of the current study are detailed below.

### 4.1. Brain Age Measures in the AUD Dataset

As shown in **Fig. 2**, the corrected brain age for the AUD individuals was 1.70 years higher than the chronological age. A vast majority of the AUD individuals, as illustrated in the distribution plot in **Fig.1**, consistently showed increased brain age. Further, there was a significant correlation between chronological age and corrected brain age (r=0.9691; **Table 3**). These findings indicate that AUD individuals, despite their current abstinence status, have accelerated brain aging. Previous neuroimaging studies have suggested that chronic alcohol use can lead to accelerated aging [21,65], as evidenced by anomalies in brain morphology as well as neurocognitive functions [66,67], which might further contribute to age-related dementia as an end outcome [21]. As the current study has used brain morphological features to estimate brain age, it is also worth mentioning that the same set of AUD participants had shown significantly smaller volumes in frontal lobe regions, such as left pars orbitalis, right medial orbitofrontal, right caudal middle frontal cortices, and in bilateral hippocampal regions in our previous study [39]. Numerous studies have confirmed that both gray and white matter volume loss in the brain contribute to accelerated aging in individuals with chronic alcohol use [68,69]. Past research has also reported that decreased cortical thickness seen in abstinent AUD individuals could be associated with severity of past alcohol misuse [70]. Furthermore, age-related cognitive decline is further accelerated by chronic alcohol misuse [71].

It may also be worthwhile to interpret the current findings in terms of other relevant domains of the brain and cognition. It is well-known that brain structural abnormalities often seen in AUD individuals are often associated with accompanying cognitive impairments [72–74], which can lead to accelerated cognitive aging and even dementia in chronic alcohol users [75–78]. This view is further confirmed by our previous findings on this dataset that the AUD participants showed relatively poorer performance in neuropsychological tests of problem-solving ability, visuospatial memory span, and working memory, compared to the healthy controls, in addition to changes in brain volume and white matter integrity [39]. Additionally, our past study also found that neuropsychological performance was correlated with several features of brain volume and white matter integrity [39], further validating the intricate relationship among brain structure, cognitive functioning, and brain aging.

### 4.2. Brain Age Measures in the FHD Dataset

The results showed that the corrected brain age for the High-FHD individuals was found to have increased by 1.08 months relative to their chronological age (**Fig. 4**). Similar to the AUD group, there was a significant positive correlation of chronological age and brain age (r=0.9875; **Table 4**), indicating that brain age increase was in tandem with chronological age. Interestingly, as illustrated in **Fig. 3**, the brain age starts increasing relative to the chronological age only after the age of 22, and the corrected brain age gap for this subset (age 22 and over, N=30) rose to 0.2334 years (2.80 months), showing that impairments in cognition and brain health may have started to appear as early as early 20s even through their clinical profiles on alcohol and substance use were not statistically different than the Low-FHD group. Previous studies have shown that children, adolescents and young adults with strong family history of AUD had reduced cortical thickness and decreased brain volume [79] and decreased brain volumes of cortical and subcortical structures [80–85]. These findings were present in alcohol-naïve high-risk offspring individuals [86], of whom a majority were yet to develop AUD or SUD [81–83,85], suggesting that these neuroanatomical differences may predate the development of AUD and reflect an underlying genetic susceptibility [87]. It is likely that genetic factors due to family history, in interaction with environmental factors, may predispose these individuals with atypical brain development [84]. Further, studies have also confirmed that these structural anomalies are also related to higher prevalence of externalizing traits, such as impulsivity and substance use behaviors, in these high-risk individuals [79,88,89], suggesting that both behavioral [90,91] and neural disinhibition [92] may predispose or mediate these individuals in developing substance use and other externalizing disorders [90–93].

Nevertheless, in the current study, it should be noted that the clinical profile (i.e., AUD/SUD diagnosis and symptom counts) between the High- and Low-FHD groups were not statistically significant, possibly suggesting that either the high-risk participants have not yet developed more severe symptoms relative to their low-risk counterparts, or alternatively, there could have been a selection bias while recruiting the participants leading to the selection of these individuals. It is quite likely that the young adults with fewer substance use problems were more readily available and amenable for the MRI scanning (especially since there is a minimum of 5- day non-drinks days required for scanning). In any case, future large-scale studies are warranted to further understand this phenomenon. However, our finding that there was a significant association between neuroanatomical features and brain age in the High-FHD group, but not in the Low-FHD individuals, is very compelling and in line with the findings from the literature. Future studies may also examine the association between brain age and neuroanatomical as well as multimodal features (genomic, clinical, and environmental factors) in different age cohorts (e.g., children, adolescents, young adults, etc.) to improve our understanding of mechanisms and interplay of various etiological factors underlying AUD and related disorders across the lifespan and brain development.

### 4.3. Brain Age and Neuropsychological Performance

Correlational analysis (Tables 6 **&** 7) showed that increase in the brain age values were significantly associated with poor scores in specific neuropsychological performance related to executive functions such as planning and problem solving in both AUD and High-FHD groups. This finding supports the view that brain age could be a reliable measure to estimate cognitive decline in various neurocognitive disorders, including in individuals diagnosed with AUD and in high-risk individuals who are likely to develop AUD later in life. Interestingly, increased brain age in the AUD group was associated with lack of efficiency in planning and problem solving (as reflected by the number of trials taken by the individuals to solve the puzzles), while on the other hand the brain age related cognitive impairment in the High-FHD group was observed in the time taken during planning and problem solving (as indicated by the pickup time and total time). These differences in cognitive profiles between AUD and High-FHD could be related to specific developmental stage and aging process of the two groups, as the AUD group is significantly older than the High-FHD group. On the other hand, in the memory task, only the chronological age showed negative association with the short-term memory performance (forward recall), suggesting poor memory performance with advancing chronological age in the AUD group. It should also be noted that the negative correlation between the same short-term memory variables and brain age was >0.39 in the AUD group, although this association did not reach statistical significance due to relatively a smaller sample size. Thus, brain age measure can be an efficient marker of cognitive performance and brain health in AUD individuals.

Overall, it is clear from our findings that increased chronological age as well as brain age were significantly associated with poor cognitive performance in both AUD and High-FHD groups, although the pattern of deficit may vary between the groups. It has been widely reported that individuals with chronic AUD diagnosis often manifest neuropsychological impairments in multiple domains, such as deficits in executive functioning, memory, and visuospatial processing [94–100]. A meta-analysis confirmed that planning, problem solving, and inhibitory abilities are significantly affected by alcohol misuse [101]. While the recovery of some cognitive processes are known to occur due to abstinence in AUD individuals, certain deficits can persist even after a prolonged abstinence [94]. In the similar vein, there is also a strong literature support to show similar neuropsychological impairments in high-risk offspring of AUD individuals. Several studies have reported neuropsychological deficits in individuals with positive family history of AUD [79,102–104]. While the High-FHD individuals, with a relatively smaller increase in their brain age, have manifested specific neuropsychological deficits in executive functioning, it is expected that these anomalies will further exasperate when they develop more alcohol problems during their mid- and later life as they advance in their age. Brain age measures could thus serve as a reliable proxy for measuring brain health and cognitive functioning in the high-risk individuals as well, as they did for the AUD individuals in this study.

### 4.4. Limitations of the Current Study and Suggestions for Future Research

While the study has yielded some interesting findings, there are some limitations that need to be mentioned. First, the sample sizes of the datasets were relatively smaller for the predictive models of machine learning, and therefore the validity of the prediction as well as the generalizability of the findings are rather limited in scope. Second, the study groups were not matched for age in the AUD dataset, in which the control participants were significantly younger than their AUD counterparts. Third, the AUD dataset comprised only male participants and therefore generalizability is limited. Fourth, in the FHD dataset, the sample sizes were not equal between the Hi-FHD and Lo-FHD groups. Fifth, the sex composition in the Low-FHD group was not perfectly matched. Lastly, while it is well-known that the diagnosis and symptoms of AUD/SUD were more prevalent in the high-risk group with a dense FHD, the study sample of Hi- FHD group did not differ from the Lo-FHD individuals in terms of their substance use profiles as seen in **Table 2**.

Future studies should use large samples of AUD and high-risk groups, preferably matched for age, sex, and other important sample characteristics to obtain realistic brain measures.

Further, the association among brain age, neural features, and behavioral/cognitive features may be examined by the future studies using samples from different age cohorts (e.g., children, adolescents, young adults, etc.) to understand potential etiological mechanisms underlying alcohol use and related disorders across brain development. Future research that would aim to study brain age in AUD related groups may also consider using other important structural and functional brain features, including brain connectivity measures, as done by some of the recent brain age studies on other neuropsychiatric disorders. Since there is a critical paucity of EEG studies on brain age in AUD and high-risk individuals, more such studies are warranted using large samples to understand subtle and nuanced brain dynamics in milliseconds time scale. It may be interesting and important to compare various prediction algorithms of both machine learning and linear models to ascertain and validate the findings across multiple studies. Finally, future studies may also try to understand correlations between brain age and measures from various domains such as neuroimaging (i.e., structural and functional MRI measures), neurocognition (e.g., neuropsychological scores), personality (e.g., internalizing and externalizing characteristics), genomics (e.g., polygenic scores), and environment (e.g., SES, family, peers, etc.).

## 5. Conclusions

Brain age measures are assuming greater importance to predict cognitive decline and accelerated biological aging in various disorders, including substance use disorders. The current study estimated brain age measures using the XGBoost regression algorithm in two distinct datasets, namely, the AUD dataset containing 60 adult participants (30 individuals with a history of AUD and 30 control individuals without any AUD diagnosis) and the FHD dataset with 71 young adults (40 individuals with high FHD scores and 31 individuals with low FHD scores). The major finding was that both AUD group and the High-FHD group showed an increase of 1.70 and 0.09 years (1.08 months), respectively, in their brain age relative to their chronological age, suggesting an accelerated brain aging and cognitive decline/deficit in these individuals. Brain age in both groups (AUD and High-FHD) was significantly associated with specific neuropsychological deficits. The accelerated brain aging in the AUD group, despite maintaining abstinence, could be due to brain damage caused by chronic alcohol use in the past. On the other hand, a small increase in brain age, relative to the chronological age, in the high FHD group may indicate an underlying neurocognitive deficit, possibly due to genetic and/or lifestyle related factors. Further studies using larger samples in different age cohorts and involving multimodal measures are needed to understand the neurobiological and genetic mechanisms underlying alcohol use and related disorders. It is recommended that future studies may circumvent the limitations of the current study and improve their research designs by using suggestions rendered from this study.

## Author Contributions

Conceptualization: C.K., B.P.; Methodology: C.K., B.P., B.A.A.; Data Collection: B.A.A.; Data Curation: B.A.A., C.K., A.K.P.; Formal Analysis: C.K., B.A.A.; Manuscript Preparation: C.K.,

B.P.; Review & Editing: B.P., B.A.A., A.K.P., J.L.M., D.B.C., C.R., S.S.d-V., W.K.; Funding Acquisition: B.P. All authors have read and agreed to the published version of the manuscript.

## Institutional Review Board Statement

The study was conducted according to the guidelines of the Declaration of Helsinki and approved by the Institutional Review Boards of SUNY Downstate Health Sciences University and Nathan Kline Institute (IRB approval ID: SUNY–266893; NKI–212263).

## Informed Consent Statement

Informed consent was obtained from all subjects involved in the study.

## Acknowledgements

In memory of Henri Begleiter, founder and longtime mentor of the Neurodynamics Laboratory, we acknowledge with great admiration his seminal scientific contributions to the field. We are sincerely indebted to his charismatic leadership and luminous guidance, truly inspired by his scientific mission and vision, and highly motivated to carry forward the work he fondly cherished. We are grateful for the valuable technical assistance of Carlene Haynes, Joyce Alonzia, Chamion Thomas, Alec Musial, Kristina Horne, and Luke Landry.

**Appendix Table A1:** List of 187 anatomical features used in the prediction of brain age.

| Cortical Thickness Features (75 LH and 75 RH ROIs) |  | Brain Volume (37 ROIs) |
| --- | --- | --- |
| 1. G&S_frontomargin | 39. Lat_Fis-ant-Horizont | 1. Left-Lateral-Ventricle |
| 2. G&S_occipital_inf | 40. Lat_Fis-ant-Vertical | 2. Left-Inf-Lat-Vent |
| 3. G&S_paracentral | 41. Lat_Fis-post | 3. Left-Cerebellum-White-Matter |
| 4. G&S_subcentral | 42. Pole_occipital | 4. Left-Cerebellum-Cortex |
| 5. G&S_transv_frontopol | 43. Pole_temporal | 5. Left-Thalamus-Proper |
| 6. G&S_cingul-Ant | 44. S_calcarine | 6. Left-Caudate |
| 7. G&S_cingul-Mid-Ant | 45. S_central | 7. Left-Putamen |
| 8. G&S_cingul-Mid-Post | 46. S_cingul-Marginalis | 8. Left-Pallidum |
| 9. G_cingul-Post-dorsal | 47. S_circular_insula_ant | 9. 3rd-Ventricle |
| 10. G_cingul-Post-ventral | 48. S_circular_insula_inf | 10. 4th-Ventricle |
| 11. G_cuneus | 49. S_circular_insula_sup | 11. Brain-Stem |
| 12. G_front_inf-Opercular | 50. S_collat_transv_ant | 12. Left-Hippocampus |
| 13. G_front_inf-Orbital | 51. S_collat_transv_post | 13. Left-Amygdala |
| 14. G_front_inf-Triangul | 52. S_front_inf | 14. CSF |
| 15. G_front_middle | 53. S_front_middle | 15. Left-Accumbens-area |
| 16. G_front_sup | 54. S_front_sup | 16. Left-VentralDC |
| 17. G_Ins_Ig&S_cent_ins | 55. S_interm_prim-Jensen | 17. Left-vessel |
| 18. G_insular_short | 56. S_intrapariet&P_trans | 18. Left-choroid-plexus |
| 19. G_occipital_middle | 57. S_oc_middle&Lunatus | 19. Right-Lateral-Ventricle |
| 20. G_occipital_sup | 58. S_oc_sup&transversal | 20. Right-Inf-Lat-Vent |
| 21. G_oc-temp_lat-fusifor | 59. S_occipital_ant | 21. Right-Cerebellum-White-Matter |
| 22. G_oc-temp_med-Lingual | 60. S_oc-temp_lat | 22. Right-Cerebellum-Cortex |
| 23. G_oc-temp_med-Parahip | 61. S_oc-temp_med&Lingual | 23. Right-Thalamus-Proper |
| 24. G_orbital | 62. S_orbital_lateral | 24. Right-Caudate |
| 25. G_pariet_inf-Angular | 63. S_orbital_med-olfact | 25. Right-Putamen |
| 26. G_pariet_inf-Supramar | 64. S_orbital-H_Shaped | 26. Right-Pallidum |
| 27. G_parietal_sup | 65. S_parieto_occipital | 27. Right-Hippocampus |
| 28. G_postcentral | 66. S_pericallosal | 28. Right-Amygdala |
| 29. G_precentral | 67. S_postcentral | 29. Right-Accumbens-area |
| 30. G_precuneus | 68. S_precentral-inf-part | 30. Right-VentralDC |
| 31. G_rectus | 69. S_precentral-sup-part | 31. Right-vessel |
| 32. G_subcallosal | 70. S_suborbital | 32. Right-choroid-plexus |
| 33. G_temp_sup-G_T_transv | 71. S_subparietal | 33. SubCortGrayVol |
| 34. G_temp_sup-Lateral | 72. S_temporal_inf | 34. TotalGrayVol |
| 35. G_temp_sup-Plan_polar | 73. S_temporal_sup | 35. SupraTentorialVol |
| 36. G_temp_sup-Plan_tempo | 74. S_temporal_transverse | 36. SupraTentorialVolNotVent |
| 37. G_temporal_inf | 75. MeanThickness | 37. EstimatedTotalIntraCranialVol |
| 38. G_temporal_middle |  |  |

